# Geography, not host identity, shapes bacterial community in reindeer lichens

**DOI:** 10.1101/2021.01.30.428927

**Authors:** Marta Alonso-García, Juan Carlos Villarreal A.

## Abstract

**Background and Aims:** Tremendous progress have been recently achieved in host-microbe research, however, there is still a surprising lack of knowledge in many taxa. Despite its dominance and crucial role in boreal forest, reindeer lichens have until now received little attention. We characterize, for the first time, the bacterial community of four species of reindeer lichens from Eastern North America’s boreal forests. We analysed the effect of two factors (host-identity and geography) in the bacterial community composition, we verified the presence of a common core bacteriota and identified the most abundant core taxa.

**Methods:** Morphological and molecular lichen species delimitation was performed based on the ITS region. The bacterial community of around 200 lichen samples was characterised using the 16S rRNA gene.

**Key Results:** Our results showed that host-lichen identity does not determine bacterial community composition in reindeer lichens, but we confirmed the influence of geography in shaping the diversity and abundance of bacteria associated to the species *Cladonia stellaris* from lichen woodlands. We also revealed that reindeer lichens share a reduced common core bacteriota composed exclusively by Proteobacteria.

**Conclusions:** The bacterial community in reindeer lichens is not host-selective. Northern lichen woodlands exhibit a significant higher diversity and abundance of bacteria associated to *Cladonia stellaris*. Nevertheless, the specific role of those bacteria as well as the process of host colonization remains to be determined. Elucidating these two aspects would be key to have a better understanding of the whole boreal ecosystems. The reduced and not diverse core bacteriota of reindeer lichens might be due to the larger size of our study area. The presence of the species *Methylorosula polaris* in the core bacteriota is evident and might have a particular importance for reindeer lichens.

## INTRODUCTION

Genomic-based microbiome research has become a popular topic over the last decade and has generated a great interest not only for the scientific community, but also for the general public. Microbiome research started in human medical studies (Turnbaugh *et al*., 2007; Lloyd-Price *et al*., 2017). Nowadays, this field of knowledge is applied to numerous domains (e.g.: vertebrate (Youngblut *et al*., 2019), insect (Douglas, 2018), plants (Trivedi *et al*., 2020), phages (Federici *et al*., 2020), soil (Delgado-Baquerizo *et al*., 2018)). As a result of this rapid interest on microbiome research, Berg et al. suggested rules and a baseline for microbiome studies, clearly delimitating the terms microbiota and microbiome (Berg *et al*., 2020). The latter contains the microbiota (community of microorganisms) and their structural elements, metabolites and the surrounding environmental conditions (Berg *et al*., 2020). Microbiome research, therefore, focuses on the interactions of microbes within a specified environment or host (Cullen *et al*., 2020). One of the most pressing questions in microbiome research is, in fact, whether exists host specificity of the microbial community. Specificity can be considered as an interaction between microorganisms and host in which absolute exclusiveness is expressed (Bubrick *et al*., 1985). It should not be mistaken with host-selectivity which describes a situation where microorganisms and host interact preferentially with one another (Bubrick *et al*., 1985). Microorganisms display various levels of host specificity, infecting a wide range of hosts (Rahme *et al*., 2000; Chappell and Rausher, 2016) or having strict host selectivity as happened in several living being (e.g., sponges (Reveillaud *et al*., 2014), hornworts (Bouchard *et al*., 2020), cetacean (Denison *et al*., 2020), humans (Pan *et al*., 2014)). Despite the impact of host identity shaping the structure and composition of microbial community, many other biotic or abiotic factors can determine the microbiota. Among the abiotic factors, geography and environmental conditions are probably the best studied (Rothschild *et al*., 2018; Zheng and Gong, 2019; Sepulveda and Moeller, 2020). Regarding host-related factors, physiology (Reveillaud *et al*., 2014; Denison *et al*., 2020), morphology (Pearce *et al*., 2017; Morrissey *et al*., 2019) or genetic (Wagner *et al*., 2016) are those who have, so far, received more attention. Random colonization and microbial interactions (Hassani *et al*., 2018) also contribute to community structure. In addition, these driving factors can interact together to determine the microbiome (Agler *et al*., 2016), and their influence can vary depending on the hosts, or the environmental conditions (Schlechter *et al*., 2019). Within the same host, each group of microorganisms may be affected by a different factor (Cardinale, Grube, *et al*., 2012), or eventually, host-individual variation in microbiome composition occurs, including individuals harboring specific taxa (Ley *et al*., 2008) that result from factors such as diet, environment, season and host physiology.

Another key aspect to considered in microbiome research is the prevalence and frequency of microorganisms in the host, namely, the core microbiome. The core microbiome was defined as a group of microbial taxa that occur with hosts above a particular occupancy frequency threshold (Risely, 2020), often between 30% (Ainsworth *et al*., 2015) and 95% (Huse *et al*., 2012) (common core). The major motivation for identifying a universal common core is to find a component of the microbiome that, due to the higher prevalence, may have a particular positive effect on the host. For example, Lee et al. detected a maintained core microbiome across jellyfish life stages which might contribute to their evolutionary success (Lee *et al*., 2018). Ainsworth et al. suggested that symbiotic core bacteria found around the endosymbiotic dinoflagellates fulfil a role in the physiology and energy requirements of coral hosts (Ainsworth *et al*., 2015). Those microbe-host symbiotic interactions have been long time studied (McFall-Ngai, 2008; Relman, 2008). Nonetheless, there are still substantial knowledge gaps in understanding the function of a core microbiome, the interactions with the host and the environment. It is universally recognized that most organisms form a symbiotic assemblage of organisms working together, otherwise known as the holobiont. The introduction of the term holobiont (Margulis, 1991) favoured the studies and promoted a holistic view on symbiotic interactions where several species are considered (Vandenkoornhuyse *et al*., 2015; Faure *et al*., 2018; Hassani *et al*., 2018; Simon *et al*., 2019). Among non-model organisms, lichens are the symbiotic organism “par excellence”, because of a partnership between fungi (one-several species), green algae, cyanobacteria and numerous bacteria in a multi-species symbiosis (Aschenbrenner *et al*., 2016; Lavoie *et al*., 2020). While the interactions between algae/cyanobacteria have been intensively studied, the role of bacteria in the symbiosis is still in its infancy (Grube *et al*., 2015). Genomic exploration of lichen associated microbes has revealed an unexpected diversity of bacteria, the majority belonging to Alphaproteobacteria (Cardinale *et al*., 2008; Grube and Berg, 2009; Hodkinson and Lutzoni, 2009; Bates *et al*., 2011; Printzen *et al*., 2012). Lichen bacteriota contribute to essential functions of host (nutrient supply, resistance against biotic and abiotic factors, growth support, detoxification of metabolites or provision of vitamin B_12_) (Grube *et al*., 2015) and, can be determine by different factors such as host-identity (Bates *et al*., 2011; Sierra *et al*., 2020), photoautotrophic symbiont (Hodkinson *et al*., 2012), thallus conditions (Mushegian *et al*., 2011; Cardinale, Steinová, *et al*., 2012; Noh *et al*., 2020) and growth form (Park *et al*., 2016), substrate type (Park *et al*., 2016), habitat (Cardinale, Grube, *et al*., 2012), or geography (Hodkinson and Lutzoni, 2009; Aschenbrenner *et al*., 2014). To date, lichen microbiomes studies have been mainly carried out in *Lobaria pulmonaria* (Cardinale, Grube, *et al*., 2012; Aschenbrenner *et al*., 2014), *Cetraria acuelata* (Printzen *et al*., 2012), *Cladonia arbuscula* (Cardinale *et al*., 2008) or *C. squamosa* (Noh *et al*., 2020). However, there is still a large and unexplored microbial diversity in other groups of lichens, such as those from northern ecosystems.

The boreal forest is the largest biome of North America. It covers 60% of the Canadian territory (Roi, 2018), extending across the continent from Newfoundland to Alaska. It stores up to 20% of global soil organic carbon (C) (Jobbágy and Jackson, 2000; Tarnocai *et al*., 2009), houses a significant number of endangered species, and is likewise crucial for indigenous human populations that have lived there for millennia (*ACIA Impacts of a Warming Arctic* 2004; Larsen, 2014). Reindeer lichens (Ahti, 1961) are terricolous lichens that have adapted better than almost all other lichens to boreal forest (Athukorala *et al*., 2016). Species such as *Cladonia mitis, C. rangiferina, C. stellaris* and *C. stygia* have become essential components of those ecosystems and, in winter, they represent the most important food source for reindeer (*Rangifer tarandus*) and caribou (*R. tarandus caribou*) (Skogland, 1984; Svihus and Holand, 2000; Thompson *et al*., 2015). In addition, they contain about 20% of the total lichen woodland (LW) biomass and can contribute up to 97% of ground cover (Auclair and Rencz, 1982; Morneau and Payette, 1989; Shaver and Chapin, 1991). Within the boreal biome, lichens are particularly dominant in LWs, a belt between the closed-canopy boreal forest to the south, and the forest tundra to the north, mostly above the 50 parallel (Payette, 1992; Johnson and Miyanishi, 1999). In Eastern North America, a remnant of LW is located 500 km south of its usual distribution range, in the *Parcs des Grands-Jardins* (PNGJ) (Jasinski and Payette, 2005).

Based on the importance of lichens in northern ecosystems as well as their utility as multi-model species to study symbiosis relationships, here we investigate the bacteriota of four lichen species to elucidate the host-microbe interactions in the boreal forest. More specifically, we (i) test host-selectivity of the bacterial microbiota associated to reindeer lichens, (ii) asses the influence of geography in composition and structure of the bacterial community of *Cladonia stellaris* from LWs and (iii) verifier the presence of a common core bacteriota in reindeer lichens. To achieve our goals, we used four lichen species (*C. miti, C. rangiferina, C. stellaris* and *C. stygia*). Bacterial host-selectivity was studied based on morphological and molecular species delimitation. The influence of geography on the bacteriota was carried out including exclusively a single species (*C. stellaris*) and a single ecosystem (LW) to reduce the bias. The presence of a southern LW in Eastern North America, makes this region an ideal setting to explore the effect of geography. Finally, we probed for bacterial taxa occurring with reindeer lichens above a particular occupancy frequency threshold and, whose presence could be interesting for the host-bacteria symbiotic interaction. This is the first extensive study of the diversity and structure of the bacterial community of the boreal forest and their interactions with host lichens.

## MATERIALS AND METHODS

### Sample collections and processing

We studied four species of reindeer lichens, *Cladonia mitis, C. rangiferina, C. stellaris* and *C. stygia* (Ahti, 1961). We collected samples along a latitudinal gradient in Eastern North America (province of Quebec) (Fig. 1). We gathered samples from the arctic tundra, forest tundra, LW (Kuujjuarapik-Whapmagoostui), closed-crown forest, balsam fir-white birch and balsam fir-yellow birch forests. We included collections from the southernmost LW in North America (PNGJ) (Jasinski and Payette, 2005).

**Fig. 1.**
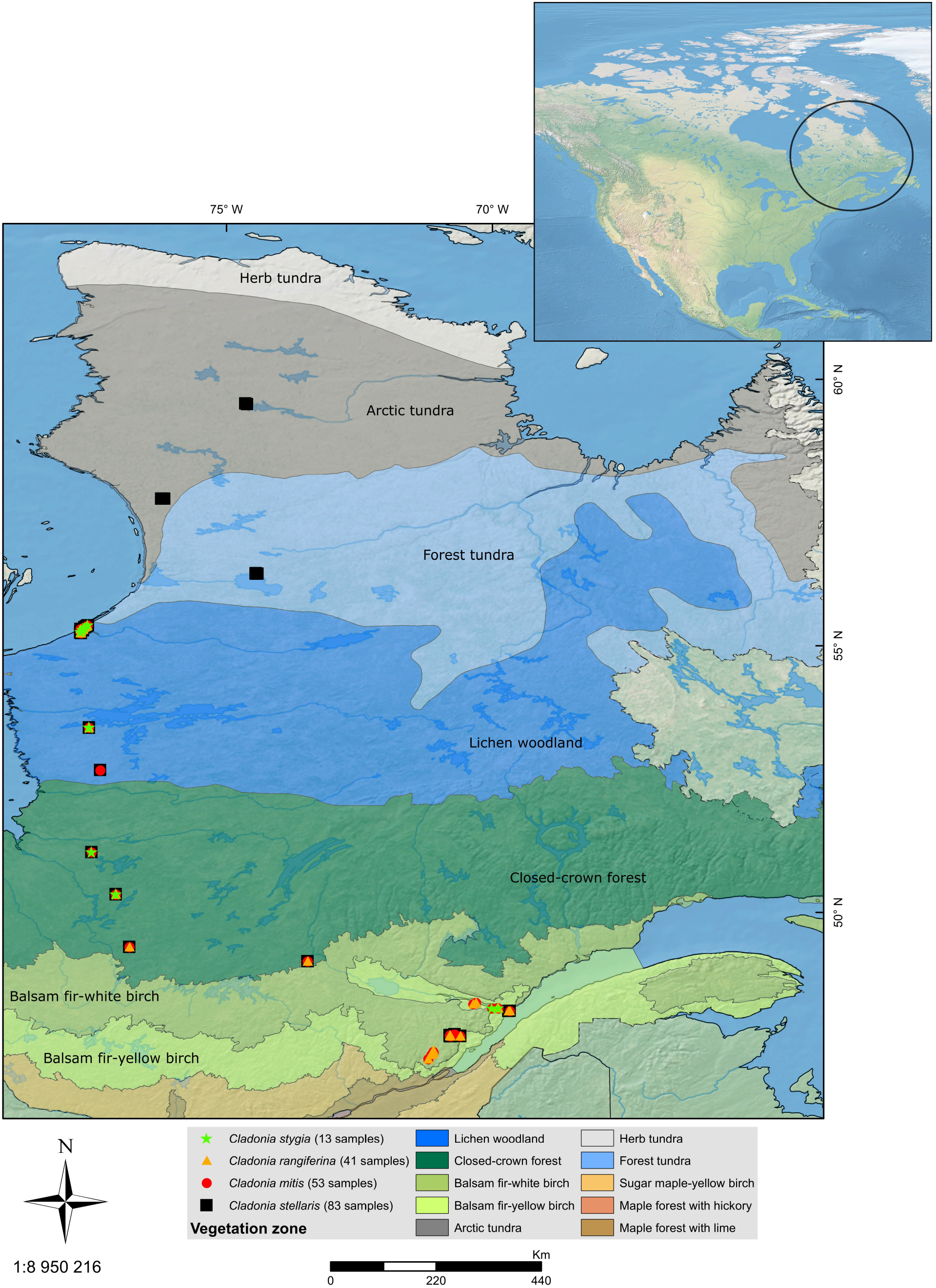
Map of Eastern North America with sampling localities of the 192 reindeer lichen specimens included in this study. Four species of *Cladonia* are represented by different symbols, red circles for *Cladonia mitis*, orange triangles for *C. rangiferina*, black square for *C. stellaris* and green stars for *C. stygia*. Bioclimatic domains highlighted following (Rowe, 1972).

Sampling tools were sterilized between collections. Samples were placed into Eppendorf tubes and stored at −20 C. Herbarium vouchers were deposited in the Louis-Marie Herbarium (QFA), Laval University. A total of 192 samples were collected and identified using regional taxonomic publications (Brodo *et al*., 2001). According to the morphological identification, 53 samples belonged to *C. mitis*, 42 to *C. rangiferina*, 84 to *C. stellaris* and 13 to *C. stygia*. Table S1 includes locality, vegetation zone (arctic, boreal or template) (Rowe, 1972), bioclimatic domain (arctic tundra, forest tundra, LW, closed-crown forest, balsam fir-white birch, balsam fir-yellow birch) (Rowe, 1972), altitude, type of genetic data generated and GenBank accession number.

### DNA extraction, PCR amplification and sequencing

Genomic DNA of lichens was extracted following an established KLC protocol (Park *et al*., 2014). The internal transcribed spacer 1 and 2 (hereafter ITS1 and ITS2) and the 5.8S of the nuclear ribosomal DNA (rDNA) were selected to perform molecular species delimitation. We successfully amplified and sequenced 104 lichen samples (forty individuals of *C. mitis*, thirty-three *C. rangiferina*, twenty-one *C. stellaris* and ten of *C. stygia*) (Table S1).

The V3-V4 region of the 16S rRNA gene was amplified following an amplicon sequencing protocol developed at Laval University (Vincent *et al*., 2017). The locus-specific primers, BactV3-V4-F (341F) and BactV3-V4-R (805R), were selected for the first PCR from (Pr Herlemann *et al*., 2011) and were modified to include Illumina TruSeq sequencing primers on their 5′ ends. PCR was conducted in a total volume of 25 μl that contained 1X Q5 buffer (New England Biolabs), 0.25 μM of each primer, 200 μM of each dNTPs, 1 U of Q5 High-Fidelity DNA polymerase (New England Biolabs), and 1 μl of DNA. The second PCR introduced indexes and Illumina adapters used in library construction. Quality of the purified PCR products was checked on a 1% agarose gel and then quantified spectrophotometrically using a NanoDrop 1000 (Thermo Fisher Scientific). The libraries were pooled using an equimolar ratio, quantified, and sequenced on an Illumina MiSeq 300-bp paired-end run at the Plateforme d’Analyses Génomiques at the Institut de Biologie Intégrative et des Systèmes (Université Laval, Québec, Canada).

### Molecular delimitation of reindeer lichen species

We conducted Bayesian inferences (BI) using the program MrBayes v.3.2 (Ronquist *et al*., 2012) on the ITS dataset, including 126 sequences (22 from GenBank), and *Cladonia wainioi* as an outgroup (Stenroos *et al*., 2018). We tested the best-fit substitution model using MrModelTest (Nylander, 2004) using the Akaike information criteria (AICc). The selected model was TrNef. Since the TrNef model is not available in MrBayes v.3.2, it was substituted with HKY+Γ (Hasegawa *et al*., 1985). The data were analysed using Markov chain Monte Carlo (MCMC), running two parallel analyses with four chains each for 20 million generations, sampling trees and parameters every 5000 generations. Chain convergence and stationarity was checked in Tracer v.1.6 (http://tree.bio.ed.ac.uk/software/tracer/), making sure the average standard deviation of split frequencies remained below 0.01, and 25% of the sampled trees were discarded as burn-in. The *allcomat* options was included to have a binary tree, required for next set of analyses. The majority consensus rule tree was visualized in FigTree v. 1.4.0 (http://tree.bio.ed.ac.uk/software/figtree/).

We tested the Poisson tree processes (PTP) model of species delimitation (Zhang *et al*., 2013) for the BI tree. PTP assumes that branching events within species will be more frequent than between species, with each substitution having a small probability of generating branching events (Kapli *et al*., 2017). Unlike other species delimitation methods such as GMYC, PTP does not require an ultrametric tree, thus eliminating potential errors and confounding effects associated with molecular dating (Dellicour and Flot, 2018; Marki *et al*., 2018). A Bayesian implementation of the PTP model (bPTP) (Zhang *et al*., 2013) was performed on the online server https://species.h-its.org/ using the BI tree. We ran 500000 generations with a thinning of 500 and a burn-in of 0.1, then assessed convergence visually using the MCMC iteration v. log-likelihood plots generated automatically.

Additionally, we applied the recently introduced multi-rate PTP (mPTP) (Kapli *et al*., 2017) method, an improved version of the PTP for single-locus species delimitation. Instead of all species sharing the same rate of evolution (λ) as in PTP, each species branch has its own λ in the mPTP model. This method determines which number of species fits best to the given data by utilizing the Akaike Information Criterion (rather than a p-value test as in PTP) because of the different number of parameters. mPTP has been shown to be consistent and very effective for species delimitation in datasets with uneven sampling (Blair and Bryson, 2017) and it has also been successfully applied to lichens (Kistenich *et al*., 2019). Using the stand-alone mPTP software (v.0.2.4) (Kapli *et al*., 2017) we performed the analyses on BI tree. We first calculated the correct minimum branch length threshold with the *--minbr_auto* option. Then, we executed a ML species delimitation inference assuming a different coalescent rate for each delimited species (*-multi}* and removing the outgroup taxa (*--outgroup crop*). To assess the confidence of the ML delimitation scheme, we conducted four MCMC runs for 20 million generations, sampling every 5000. The first two million generations were discarded as burn-in and analyses started with a random delimitation (*--mcmc_startrandom*). We compared results among MCMC runs to assess congruence.

### Taxonomic assignment of bacterial sequences

We used the DADA2 workflow to assign Amplicon Sequence Variants (ASVs; (Callahan *et al*., 2017)) in R (Callahan *et al*., 2016). We trimmed and filtered the raw reads, keeping only those with quality scores higher than 25. We dereplicated (*derepFastq*) all reads, estimated their error rates (*learnErrors*) and denoised them (*dada*). Then, forward and reverse reads were merged (*mergePairs*). All merged sequences with less than 430 bp and more than 450 bp were removed, and chimeras were excluded. We assigned taxonomy based on the SILVA 138 database (McLaren, 2020) with minimum bootstrap set to 80.

The phangorn R package (Schliep, 2011; Schliep *et al*., 2017) was used to build a phylogenetic tree, used in downstream analyses and to estimate phylogenetic distances between microbial communities. We first built a neighbor-joining tree, and then fit a GTR+Γ+I (Generalized time-reversible with gamma rate variation) maximum likelihood tree using the neighbor-joining tree as a starting point.

We synthesized all the data generated [ASV table, sample data, taxonomy table, phylogenetic tree and environmental variables (Table S1)] into a single phyloseq object with the R package phyloseq (McMurdie and Holmes, 2013). ASVs corresponding to chloroplast and mitochondria were removed from the dataset along with the bacteria for which no phylum could be assigned. We also removed phyla with less than ten corresponding ASVs in all samples combined. We determined prevalence (fraction of samples in which an ASV occurs) and used it to create a second phyloseq object including only the ASVs occurring in at least 5% of the samples.

### Characterization of bacterial communities in reindeer lichens

We applied diversity and composition analyses to two set of data, host-species dataset to test bacteria host-selectivity in reindeer lichens (objective i), and LWs dataset to assess the influence of geography in *C. stellaris* (objective ii). With the R package phyloseq (McMurdie and Holmes, 2013), we estimated alpha-diversity within each sample based on number of observed ASVs, Shannon and Simpson effective indices. Shapiro-Wilk tests indicated if the data were normally distributed. We then used parametric (ANOVA) or nonparametric (Kruskal-Wallis and U-Mann-Whitney tests) statistics methods to test for significant differences. To evaluate beta-diversity, meaning diversity between samples, the UniFrac distance matrix was calculated. Unlike other metrics, UniFrac takes into account phylogenetic information. Principal Coordinates Analysis (PCoA) and Double Principal Coordinates Analysis (DPCoA) based on the UniFrac distance matrix were plotted. To determine whether bacterial communities significantly change, PERMANOVA tests and pairwise comparison were conducted (2000 permutations) using the adonis and pairwise.adonis (Martinez Arbizu, 2020) functions in the vegan R package (Oksanen *et al*., 2020). We transformed ASV counts per sample into relative abundance and compare it among species, and LWs. To identify specific ASVs that show differential abundance among taxa, the R package DESeq2 (Love *et al*., 2014) was used. We removed samples with less than 1000 reads and applied a normalized logarithmic transformation (rlog) on the ASVs. We estimated logarithmic fold change (LFC) and dispersion for each ASV. We obtained an adjusted p-value (padj) to corrected for false positives (False Discovery Rate, FDR) using the Benjamini-Hochberg (BH) correction. These ASVs sequences were aligned to sequences in NCBI database 16S ribosomal RNA (Bacteria and Archae) using Megablast optimize for highly similar sequences. The identifications considered successful were those with over 97% similarity. Below this percentage, we treated the sequences as relatives.

### Detection of core bacterial members in reindeer lichens

To have a comprehensive estimate of the bacteria occurring across host reindeer lichens, the core bacteriota was identified with the microbiome R package (Lahti,). Due to the lack of consensus about a fixed threshold (Risely, 2020), we considered as core bacteriota any taxon with a prevalence higher than 0.50, 0.75 or 0.90 (Jorge *et al*., 2020). We included 189 lichen samples in the analysis. We pruned out the phyloseq object to retain those taxa with more than 1000 reads; in the end 153 samples were incorporated. We detected the core bacteriota with the function (*core members*) and estimated the total core abundance in each sample (*sample sums*). A heatmap was elaborated to visualize the results. In order to identify the core taxa as species level, ASVs sequences were aligned to sequences in NCBI database 16S ribosomal RNA (Bacteria and Archaea) using Megablast optimize for highly similar sequences and retaining those with Blast hits over 97% identity. Bacterial sequences with identity below 97% were considered as relatives.

Due to the predominance of *C. stellaris* over the remaining reindeer lichens in Eastern Canada, we carried out the same analyses to predict the core bacteriota of this species using both the morphological and molecularly delimitated taxa.

## RESULTS

### Identity of reindeer lichen species

We successfully amplified the ITS locus of 104 samples representing four morphological species of *Cladonia, C. mitis, C. rangiferina, C. stellaris* and *C. stygia*, and 22 sequences from GenBank. The alignment had a total length of 906 bases (Appendix 1). Five hundred seventy-nine sites were conserved, 321 variable and 229 parsimony informative. The BI phylogenetic tree supported (PP ≥ 0.95) a clade including all the reindeer lichens (Fig. 2). *Cladonia stellaris* split into two groups, an unsupported (Clade *C stellaris - species 2*, Fig. 2) and a supported clade (Clade *C stellaris - species 1*, Fig. 2), with the only exception of a single specimen from GenBank (accession number KP001212). Specimens of *C. mitis, C. rangiferina* and *C. stygia* clustered in a supported clade split into six unsupported subclades, four including *C. mitis* and two grouping *C. rangiferina* and *C. stygia*. The tree was used for species delimitation analyses.

**Fig. 2.**
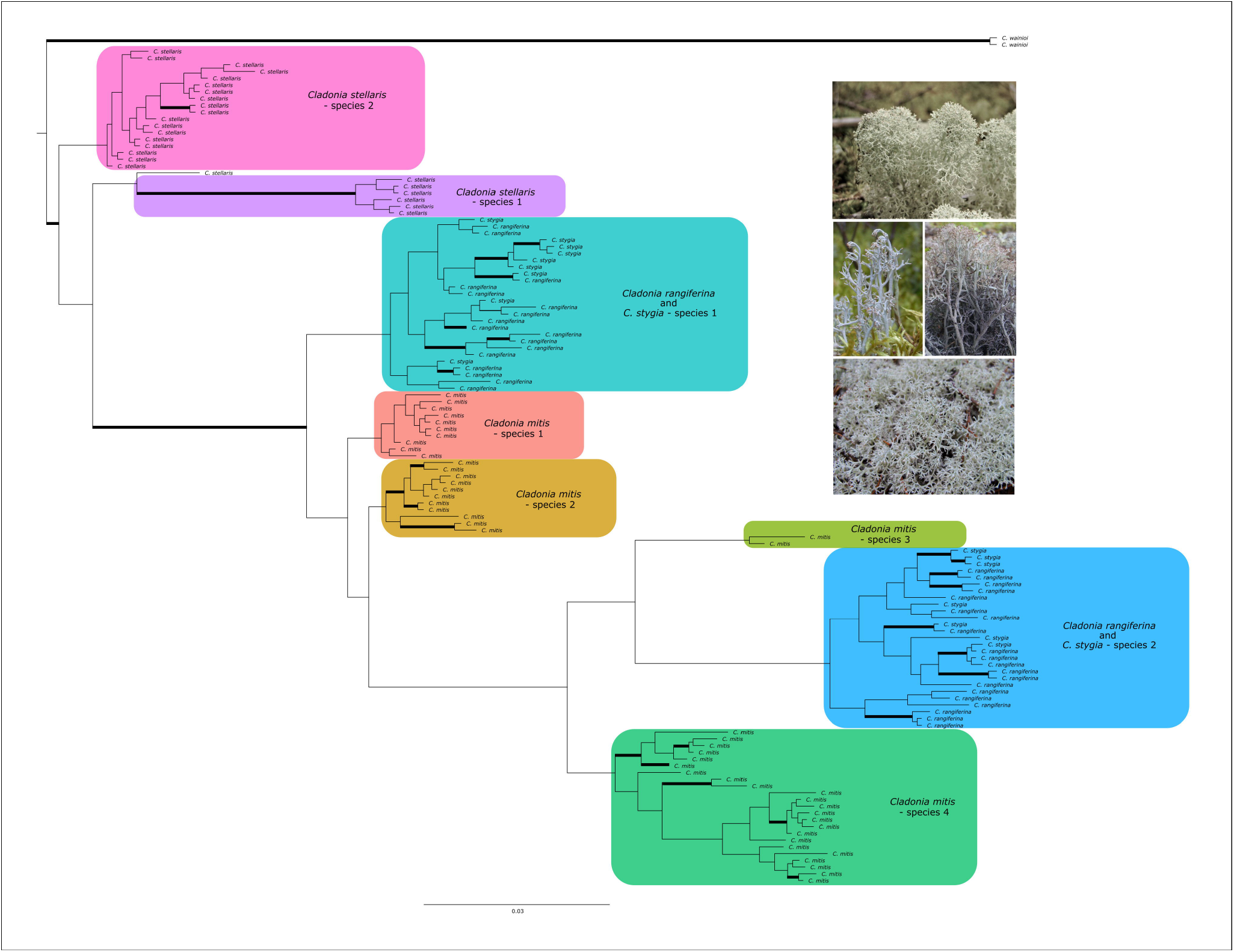
Majority-rule consensus tree of a Bayesian inference analysis from 126 ITS accessions of reindeer lichens. Branches in bold are supported by posterior probabilities (PP ≥ 0.95). Each color represents a species delimited by the multi-rate Poisson tree processes method (mPTP). Pictures show the four morphologically recognized species.

The MCMC chains for the bPTP species delimitation method did not converge (Fig. S1) thus, the model not further pursued. The mPTP method inferred ten species, with identical results for the ML and the MCMC analyses. The ML delimitation results were strongly supported (average support values over 0.91), indicating that there is a 91% congruence between the support values and the point estimates. Equally, independent MCMC runs can also be quantify using the average standard deviation of delimitation support values (ASDDSV). Here, our four independent MCMC runs converged with ASDDSV below 0.001 (Kapli *et al*., 2017). Fig. 2 shows the BI phylogenetic tree with the species identified by the mPTP method. *Cladonia mitis* was divided into four taxa; *C. stellaris* split into three taxa, one of them including exclusively a single specimen from GenBank; finally, individuals belonging to *C. rangiferina* and *C. stygia* were merged together but split into two species (Appendix 2).

### Dominance of Proteobacteria in reindeer lichens

A total of 6204737 raw reads from 16 rRNA were identified from 189 samples of reindeer lichens (Table S2). After quality trimming, 817688 reads were kept. High-quality reads were taxonomic assigned to 12917 ASVs. Once we removed chloroplast, mitochondrial and phyla with less than ten ASVs, we retained 10969. The prevalence filter (ASVs occurring in at least 5% of the samples) preserved 724 ASVs in ninety-eight samples (host-species dataset); and 1130 ASVs in forty-seven samples (LWs dataset). All the ASVs could be assigned at phylum level. The ASVs corresponded to Acidobacteriota, Cyanobacteria, Planctomycetota, Proteobacteria, Verrucomicrobiota and Eremiobacterota (candidate division WPS-2). Most of the sequences belonged to Proteobacteria (ca. 80%), followed by Eremiobacterota. Table S3 displays number of ASVs detected for each dataset at phylum level. Bacteria relative abundance of each lichen sample is showed in Fig. S2.

### Absence of bacteria host-selectivity in reindeer lichens

The bacterial alpha diversity within species delimited by mPTP methods (Fig. 3A) was significant between *C. rangiferina plus C. stygia sp 2* and *C. stellaris sp 1* in observed ASVs (p-value = 0.00084) and Shannon index (p-value = 0.00061) (Appendix 3). No significant differences in diversity were detected for the Simpson index (p-values > 0.05) (Appendix 3). In the PCoA, axis 1 and 2 explained 33.7% and 26.7% of the total variation among samples, respectively (Fig. S3A). In the DPCoA, CS1 explained 43.4% and CS2 explained 29.9% of the total variation (Fig. 4A). Bacterial communities of reindeer lichens were not grouped according to host species, and PERMANOVA tests found no significant association (Appendix 4). When considering morphological species (Fig. S4), diversity was significantly different among *C. mitis, C. rangiferina* and *C. stellaris* (p-values < 0.05), but it was not the case for *C. stygia* (Appendix 3).

**Fig. 3.**
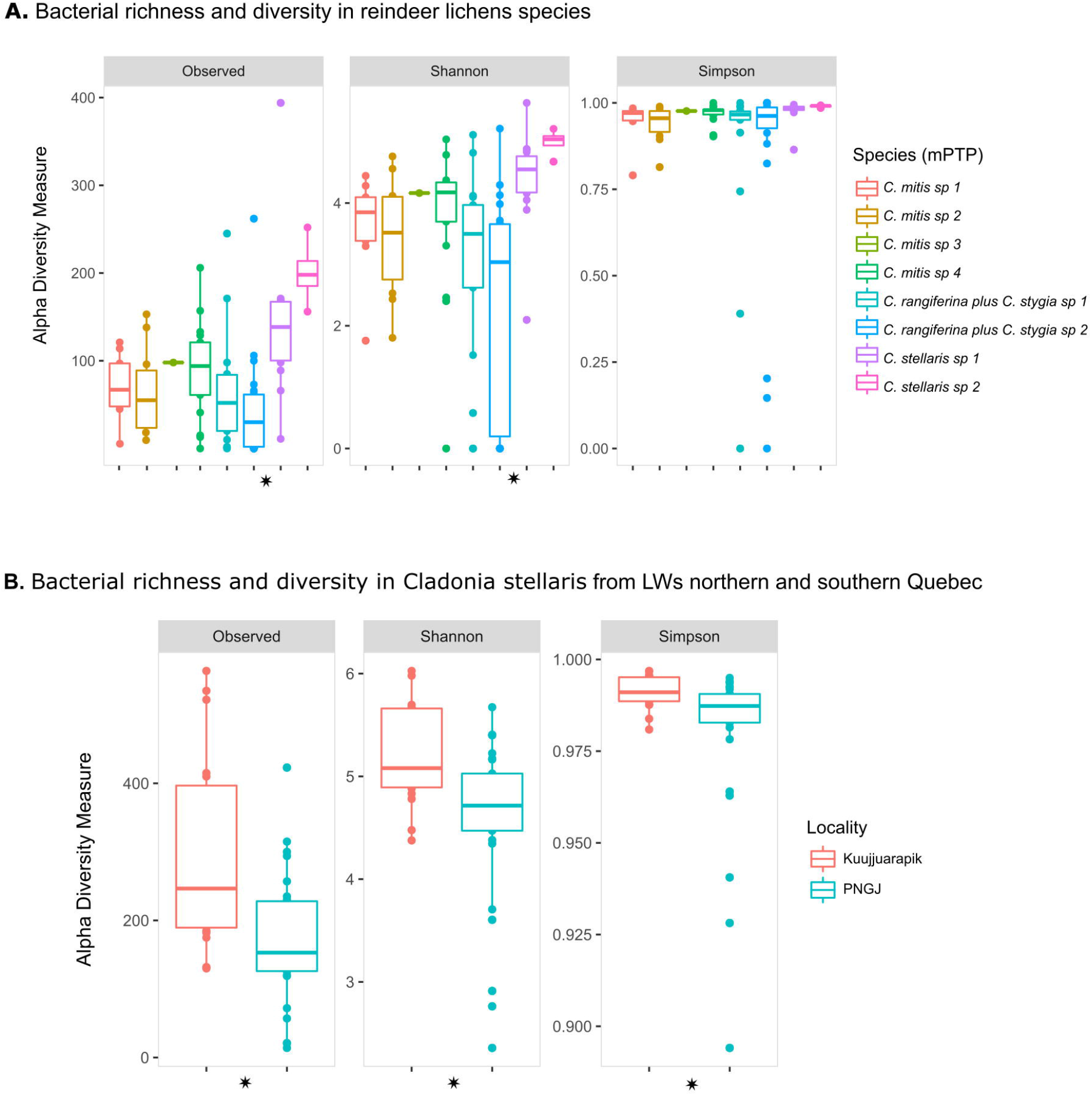
Richness estimate values (number of observed amplicon sequence variants, ASVs) and alpha diversity indices (Shannon and Simpson) for reindeer lichens bacterial communities. A. Diversity values of reindeer lichens species delimited by the multi-rate Poisson tree processes method (mPTP) (host-species dataset). The species colors follow those of Fig. 2. B. Diversity values of *Cladonia stellaris* from lichen woodlands (LWs) northern and southern Quebec (LWs dataset). Asterisks indicate significant differences (p-value < 0.01).

Proteobacteria were dominant across all reindeer lichens (Fig. S2A), although there were some exceptions. Two lichens taxa, one identified as *C. mitis sp 1 (Alonso 422*) and the other one as *C. rangiferina plus C. stygia sp 1 (Alonso 433)*, harbored mostly Eremiobacterota. Two other lichens belonging to *C. rangiferina plus C. stygia sp 2* included only Cyanobacteria (*Alonso 423*) or Planctomycetota (*Alonso 475*). DESeq2 analysis identified ASVs that displayed a significant change of abundance (padj < 0.05) compared to host lichen species (Table S4A, S4B). Bacterial communities of reindeer lichens did not exhibited differences in abundance depending on the host-lichen species. For the eight molecularly delimitated species (mPTP method), a single member of family Acetobacteraceae was detected (Table S4A), whereas for the four morphological species, one ASV belonged to Acidobacteriaceae was significantly different (Table S4B).

### Effect of geography in the bacterial community of C. stellaris

Lichens from northern and southern LWs exhibit significant differences (p-values < 0.05) in bacterial diversity (Fig. 3B) (Appendix 5). Number of observed ASVs as well as Shannon and Simpson indexes reported higher diversity for northern samples (Fig. 3B). The two first axes of the PCoA explained 28.1% and 21.8% of the total variation, respectively (Fig. S3B), whereas axes 1 and 2 of DPCoA explained 31.1% and 26.6%, respectively (Fig. 4B). Both ordination methods showed two different bacteria groups based on the latitude, northern and southern LWs (Figs. S3B, 4B), and PERMANOVA test confirmed this association (p-values < 0.05) (Appendix 6). Relative abundance analysis pointed out the higher abundance of Eremiobacterota in northern LWs (Fig. S2B), with the exception of 5 samples where this phylum was nearly absent (samples *Alonso 296, 312, 315, 326* and *354* in Fig. S2B). A total of 63 ASVs were significantly different in abundance between northern and southern LWs (Table S4C, Fig. S5A). The genus *Endobacter* (Acetobacteraceae) was the only group significantly more abundant in the south (PNGJ) than in the north (Kuujjuarapik) (Fig. S5B). According to the Blast alignment (Appendix 7), their closest known relative was *Gluconacetobacter tumulioli* with about 95.3% identity (Table S4C). The remaining ASVs were always more abundant in LWs northern Quebec (Kuujjuarapik) (Fig. S5B) and they seem to be related to *Rhodopila globiformis* (Acetobacteraceae) (96.8% identity). *Tundrisphaera lichenicola* (Isosphaeraceae) was identified using the Blast algorithm (94.9% identity) (Table S4C). The closest relative of Eremiobacterota (15 ASVs) is the bacterium species *Desulfofundulus thermocisternus*, although its percentage of identity is below 85% (Table S4C). Finally, the family Caulobacteraceae might be represented by species related to *Brevundimonas vesicularis* (96.6% identity) (Table S4C).

**Fig. 4.**
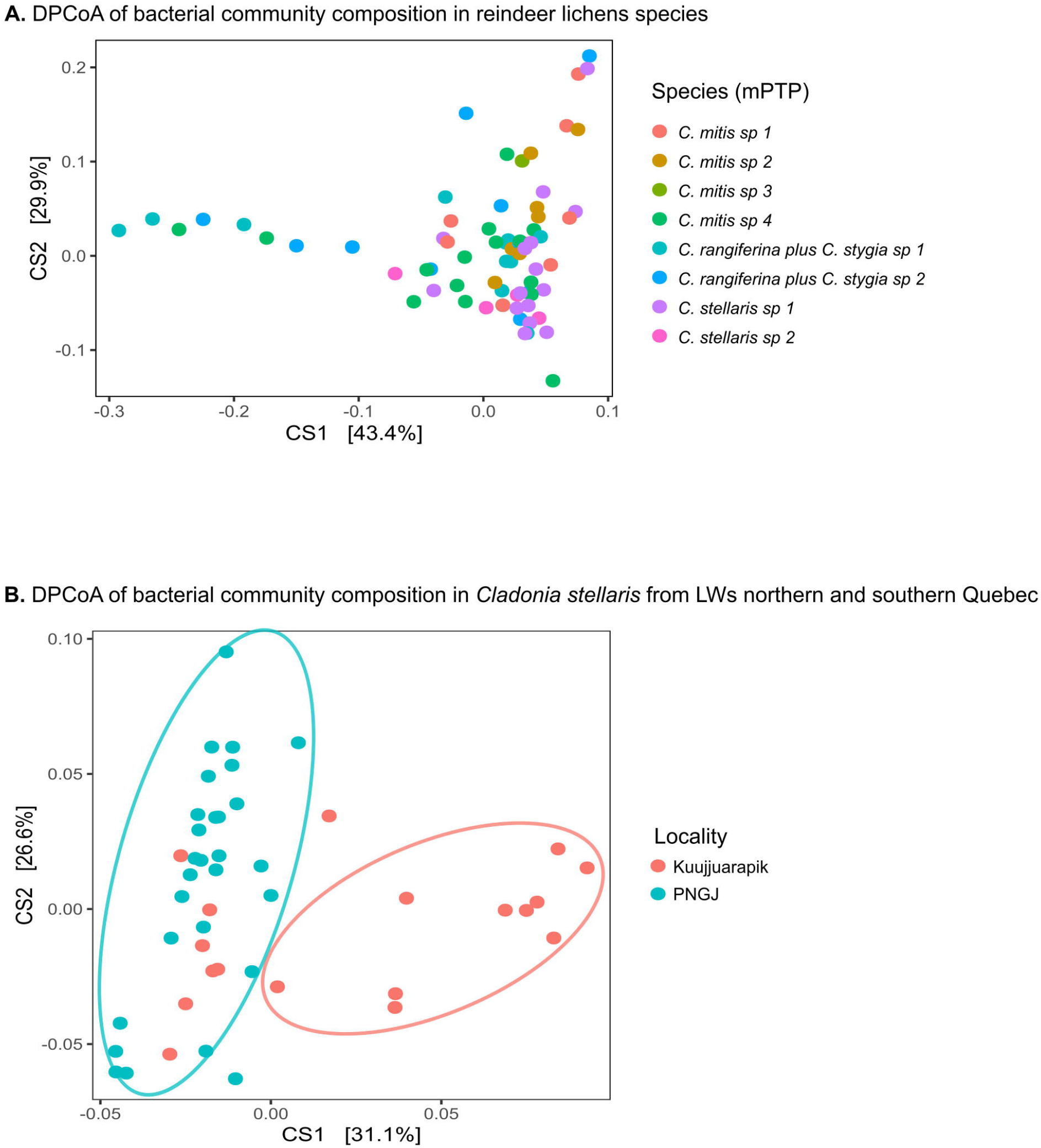
Double Principal Coordinates Analysis (DPCoA) of bacterial community composition based on amplicon sequence variants (ASVs) (A) in reindeer lichens and, (B) in *Cladonia stellaris* from lichen woodlands (LWs) northern and southern Quebec. Bacterial communities are not grouped by host species. Two clusters are differentiated between northern and southern LWs (p-value < 0.01). The species colors in (A) follow those of Fig. 2.

### Reduced core bacteriota in reindeer lichens

Forty-five ASVs were present in 153 reindeer lichens based on 0.50 prevalence-cut-off (Table S5A), representing a minor part (ca. 5%) of the total number of ASVs (863 ASVs). All of them belonged to Proteobacteria, 32 were identified as genus *Methylocella* (family Beijerinckiaceae). A single ASV from Beijerinckiaceae as well as 12 members of family Acetobacteraceae were not assigned at genus level. Using the prevalence threshold of 0.75, one ASV (*Methylocella*) was detected (Table S5B), whereas no ASV occurred in all reindeer lichens at a prevalence of 0.90. The relative abundance of the common core bacteriota was variable among lichen samples. For a prevalence of 0.50, it ranged from 2% in two samples of *C. stellaris* (Table S6) to 52% in one individual of *C. mitis* (Table S6). Using 0.75 as prevalence level, the highest relative abundance of the single ASV was 13% in one sample of *C. mitis*. Thirty-six of the 153 lichens lack this ASV (0% of relative abundance) (Table S6). The heatmap indicates the prevalence of each ASV for each abundance threshold (Fig. 5A). Four *Methylocella* ASVs presented the highest prevalence (yellow ASVs). The results derived from the Blast alignment are shown in Appendix 8. The potential identity of each ASV based on NCBI database is included in Table S5. *Methylorosula polaris* (identity over 97.7%) and sequences belonged to order Rhodospirillales (identify below 97%) were found in the common core bacteriota.

**Fig. 5.**
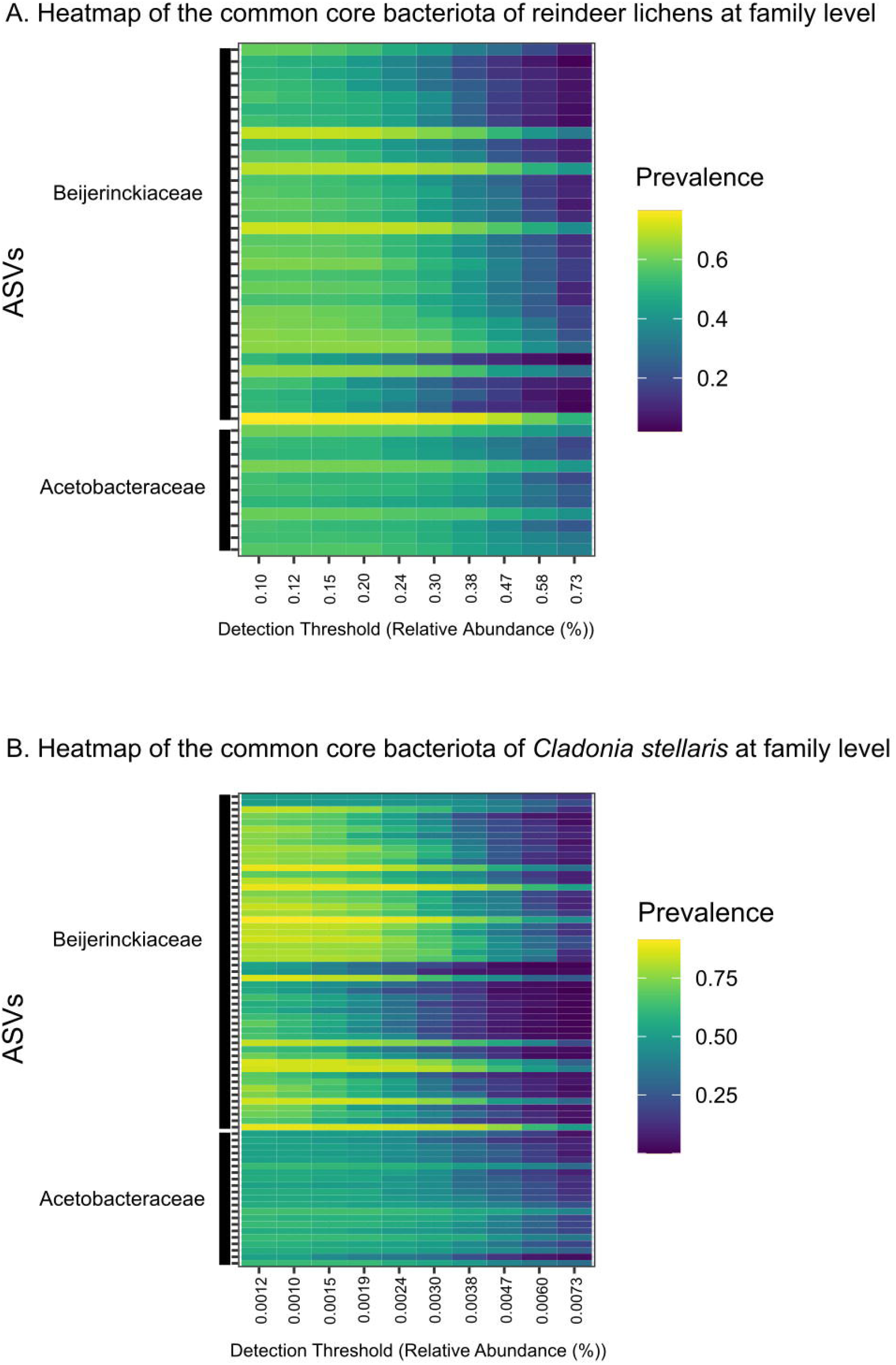
Common core bacteriota of reindeer lichens at family as a function of the abundance threshold for (A) reindeer lichens, and for (B) *Cladonia stellaris*, with prevalence above 0.50. The x-axis represents the detection thresholds (indicates as relative abundance) from lower (left) to higher (right) abundance values. Color shading indicates the prevalence of each bacterial family among samples for each abundance threshold. As we increase the detection threshold, the prevalence decreases.

A total of 81 samples of *C. stellaris* shared 87, 34 and one ASVs for prevalence of 0.50, 0.75 and 0.90, respectively (Table S7). The heatmap showed higher levels of prevalence for members of family Beijerinckiaceae than for the Acetobacteraceae (Fig. 5B). With the best Blast hits, the common core bacteriota was constituted by 55 taxa of the bacteria species *Methylorosula polaris* (identity over 97.7%) and 30 taxa closely related to *Granulibacter bethesdensis* (identity around 96.6%) (Appendix 9). Two relatives of *Methylocystis bryophila* were also included in the common core of *C. stellaris* with a percentage of identity of ca. 94%. A Venn diagrams displayed the overlap between the common core bacteriota of reindeer lichens (47 ASVs) and that of *C. stellaris* (Fig. S6). All the ASVs associated to reindeer lichens were included in the common core of *C. stellaris*.

## DISCUSSION

In this study, we present the first characterization of the bacterial community in reindeer lichens from Eastern North America. We analysed the influence of two factors (host-identity and geography) in shaping the bacterial community composition, we verified the presence of a common core bacteriota and we identified the most abundant core taxa in all reindeer lichens, with emphasis on *C. stellaris*. Our results showed no changes of the bacterial community among host reindeer lichens, but significant differences in diversity and abundance of bacteria associated to *C. stellaris* from northern or southern LWs. We also revealed that reindeer lichens share a reduced common core bacteriota composed exclusively by Proteobacteria.

### Host-lichen identity does not determine bacterial community composition

In order to test host-selectivity of the bacterial community in reindeer lichens, four morphological species were delimited based on the ITS molecular marker. This is the first attempt to compare molecular and morphological host-species delimitation in a microbiome study. Among the four morphological reindeer lichen species included here, the mPTP method identified eight species, four *C. mitis*, two *C. stellaris* and two *C. rangiferina* and *C. stygia* grouped together. The analyses of the bacterial community were applied to molecular and morphological species. Bacterial diversity within each sample hardly varies between the eight host species (delimited based on mPTP), but differences were observed between morphological species. Sierra et al. compared bacterial community in seven lichen genera, and they showed differences in alpha diversity between *Usnea* and *Hypotrachyna* (Sierra *et al*., 2020), although there was no information available at species level. Nonetheless, our results must be interpreted with caution. Single-locus species delimitation methods have limitations (Blair and Bryson, 2017; Dellicour and Flot, 2018). Here, we have used the mPTP and bPTP methods to compare different approaches, but we are aware that the molecular species delimitations might change using a different approach or increasing the number of loci.

In terms of abundance, all reindeer lichens harbour similar number of bacteria which do not cluster together by host species (neither molecular nor morphological species). Our results differ from previous studies which proved that the bacterial community in lichens were host-specific (e.g., (Grube *et al*., 2009; Bates *et al*., 2011; Sierra *et al*., 2020)). Grube et al. found species-specific bacteria in three lichens with different growth form, fruticose, crustose, and foliose (Grube *et al*., 2009). Sierra et al. also suggested bacteria host-specificity in seven genera of lichens with different thallus morphology (*Cora*, *Hypotrachyna, Peltigera* and *Sticta* are foliose, and *Usnea, Cladonia* and *Stereocaulon* are fruticose) (Sierra *et al*., 2020). We suggest that host-specificity might be, in fact, growth-form-specificity. Our study focuses only on fruticose species with similar thallus morphology, which might explain the lack of selectivity. Park et al. demonstrated that bacteria grouped together depending on the growth forms of the lichen host (crustose, foliose or fruticose) (Park *et al*., 2016). This assumption agrees with (Fernández-Brime *et al*., 2019) who revealed that morphologically simple forms of lichenization (borderline lichens) do not influence bacterial communities but, complex thallus structure are required for the lichens to provide unique niches to host specific bacterial communities. A similar pattern was observed in green seaweeds where the bacterial variation was attributed to thallus differentiation (Morrissey *et al*., 2019), or in the liverwort *Marchantia inflexa* whose differences in bacteriome seem correspond to differences in the physiology and morphology of male and female plants (Marks *et al*., 2017).

### Geography shapes the bacterial community composition

Alonso-García et al. revealed that populations of *C. stellaris* in southern Quebec were not genetically different from those of northern LWs, and they suggested constant migration of lichen individuals between populations (Alonso-García *et al*., 2021). However, the bacterial community associated to *C. stellaris* does not follow this pattern. We found significant differences between Kuujjuarapik and PNGJ’s LWs. Lichens from Kuujjuarapik (northern LWs) exhibit higher diversity and abundance of bacteria. The effect of geography in the bacterial community of other lichens had been previously tested. Hodkinson et al. used different species of lichens to show that bacterial communities were significantly correlated with differences in large-scale geography (Alaska, Costa Rica, and North Carolina) (Hodkinson *et al*., 2012). At smaller scale, the bacterial community of *L. pulmonaria* from the same sampling site showed higher similarity than those of distant populations (100 km of linear distance) (Aschenbrenner *et al*., 2014). Bacteria associated to *C. aculeata* were, likewise, affected by geography although, contrary to our results, they were less diverse in high latitude (Antarctica and Iceland) than in extrapolar habitats (Spain and Germany) (Printzen *et al*., 2012).

Differences in bacterial relative abundance between LWs were particularly evident for Eremiobacterota, a phylum usually found in acidic and cold environments (Grasby *et al*., 2013; Trexler *et al*., 2014; Bragina *et al*., 2015) and putatively capable of anoxygenic carbon fixation in boreal mosses (Holland-Moritz *et al*., 2018), although their role in reindeer lichens is unknown. Members of the family Caulobacteraceae (represented by species related to *Brevundimonas vesicularis*, ca. 96.6% identity) were also more abundance in northern LWs. This family has been frequently associated to lichens (Hodkinson and Lutzoni, 2009; Hodkinson *et al*., 2012; Aschenbrenner *et al*., 2014; Sigurbjörnsdóttir *et al*., 2015; Park *et al*., 2016; Noh *et al*., 2020), but their functions have never been elucidated. Seven ASVs closely related to the anoxygenic phototrophic purple bacterium *Rhodopila globiformis* (Acetobacteraceae) were, likewise, significantly more abundance in Kuujjuarapik. The order Rhodospirillales is dominant in Antarctic lichens (Park *et al*., 2016), and it may provide photosynthetic products, defends against pathogens and reduces the oxidative stress (Cernava *et al*., 2017). Finally, with an identity of 94.9%, we found a close relative of *Tundrisphaera lichenicola*, significant more abundance northern Quebec. This bacterium of the phylum Planctomycetes was described from lichen-dominated tundra soils within the zone of forested tundra and discontinuous permafrost of northwest Siberia (Ivanova *et al*., 2016; Kulichevskaya *et al*., 2017). To comprehend why northern LWs harbour more bacteria, we suggest focusing on the origin of those bacteria and the reason that encourage them to colonize the lichen. In addition, future studies should clarify the role that Eremiobacterota, Caulobacteraceae, Rhodospirillales and Planctomycetes play for reindeer lichens and/or for the boreal forest to better understand patterns of diversity and abundance.

Unlike aforementioned bacteria, the genus *Endobacter* (with *Gluconacetobacter tumulioli* as the closest known relative according to NCBI dataset) was more abundant in southern LWs. The specific role of *G. tumulioli* in the ecosystem remains unknown (Nishijima *et al*., 2013), although other species of this genus, such as *G. diazotrophicus*, are involved in nitrogen fixation (Saravanan *et al*., 2008). If this is the case, studies should be carried out to investigate the reason why populations northern Quebec lack these nitrogen-fixing bacteria and how they supply nitrogen shortage.

### Common core bacteriota is limited and homogeneous

We have defined a common core bacteriota as the bacteria occurring with reindeer lichen above an occupancy frequency threshold of, at least, 0.50. Comparing 153 samples throughout Eastern Canada revealed a common core of 45 ASVs representatives of families Acetobacteraceae and Beijerinckiaceae (orders Acetobacterales and Rhizobiales, respectively). All these members were also found to be associated with the single species *C. stellaris*. Sierra et al. identified a reduced (16 OTUs, threshold ≥ 0.90), but more diverse core in different genera of Paramos’ lichens (orders Rhodospirillales, Sphingomonadales, Rhizobiales, Acidobacteriales, and the phylum Cyanobacteria) (Sierra *et al*., 2020). The core bacteriota of Austrian populations of *L. pulmonaria* represented 16% of the OTUs (ca. 5% in reindeer lichens) from six phyla (Alphaproteobacteria, Sphingobacteria, Actinobacteria, Nostocophycideae, Spartobacteria and Deltaproteobacteria), but it was considered as a regional core (Aschenbrenner *et al*., 2014). The reason why reindeer lichens hardly share a core bacteriota might be due to the larger size of our study area. Detecting the core across sites or populations can provide a more reduce core bacteriota but, as suggested by (Risely, 2020), allow us to identify potential candidates for further investigation with regard to host-microbe interactions. Nevertheless, we should consider that bacteria can have high occupancy frequency within the host population for many reasons (e.g., they are common in the environment (David *et al*., 2014) or highly competitive against other microbes (Coyte and Rakoff-Nahoum, 2019)) and is not necessarily linked to host function.

Blast hits of most of the ASVs composing the core bacteriota of reindeer lichens belonged to *Methylorosula polaris* (ca. 97.93% identity), a member of the order Rhizobiales isolated from methane-oxidizing communities of soil from the polar tundra (Berestovskaya *et al*., 2012). *Methylorosula polaris* had been previously detected (identity 94.5%) at the apical and middle parts of *C. squamosa* thalli (Noh *et al*., 2020). In general, Rhizobiales occurs in lichens (Hodkinson and Lutzoni, 2009), where they are known to perform functions supporting the symbiosis, including auxin and vitamin production, nitrogen fixation and stress protection (Erlacher *et al*., 2015; Cernava *et al*., 2017). The remaining ASVs from the core bacteriota belonged to Acetobacterales and they might be relatives of the human pathogen *Granulibacter bethesdensis* (ca. 96.6% identity) (Greenberg *et al*., 2006), the phototrophic *Rhodophila globiformis* (ca. 96.4% identity), *Endobacter medicaginis* (ca. 96.1% identity) or different species of genus *Gluconacetobacter* (ca. 95.9% of identity) involved in nitrogen-fixation (Fuentes-Ramírez *et al*., 2001; Saravanan *et al*., 2008).

*Cladonia stellaris* harbours almost the same common core bacteriota than all reindeer lichens. The main different is due to the number of ASVs, which is higher in *C. stellaris* (45 versus 87 ASVs). In addition, *C. stellaris* included *Methylocystis bryophila* (Belova *et al*., 2013), a bacterium isolated from acidic *Sphagnum* peat in Europe.

### Conclusions

We provide the first assessment of the lichen microbiome in the boreal forest of Eastern Canada. Here, we answer some key aspect of lichen bacteriome from northern ecosystems and highlight future research venues. We show a dominance of Proteobacteria in reindeer lichens and an absence of bacteria host-selectivity. We suggest that thallus morphology, and consequently growth-from, may have an effect on the bacterial community composition. Our results evidence the influence of geography in shaping the bacterial community of reindeer lichens. A single species from one particular ecosystem exhibits significant higher diversity and abundance of bacteria in northern lichen woodlands. Further studies should include environmental variables, such as temperature, humidity, soil pH or light intensity, to elucidate whether abiotic factors also influence the microbial community of lichens from the boreal forest. Likewise, we suggest examining the role of soil microbial communities as a source of bacteria and the way they colonize the lichen. Regarding the core bacteriota, we identify a reduced core in reindeer lichens comprised mainly of *Methylorosula polaris*. A deeper understanding of the interaction between reindeer lichens and *Methylorosula polaris* would help to discover ecological and functional processes at the organismal and ecosystem level.

## SUPPLEMENTARY DATA

Supplementary data are available online at XXXXX and consist of the following. Six figures; seven tables; nine appendices and three scripts. The datasets generated and analysed during the current study are available in the NCBI SRA archive under Bioprojects PRJNA593044 and PRJNA687262.

## FUNDING

This research was conducted with financial support from the Spanish “Fundación Séneca-Agencia de Ciencia y Tecnología de la Región de Murcia” (project 20369/PD/17); the program Sentinel North financed by the Canada First Research Excellence Fund (CFREF); the CRNSG- RGPIN/05967-2016 and Fondation Canadienne pour I’innovation (project 36781).

## ACKNOWLEDGEMENTS

We thank the “Plateforme Analyse génomique” and “Plateforme Bio-informatique” from the “Institut de Biologie Intégrative et des Systèmes” (Laval University, Quebec) for sequencing and for advice during the analysis of data. All manipulation of sequence reads was implemented on the UNIX server of Laval University. We thank the authorities of the “Parc national des Grands-Jardins (SEPAQ)” for giving the permission to collect samples. We are also grateful to Catherine Chagnon for samples northern Quebec and Troy McMullin for pictures of lichen species.

---

Fig. S1. Markov chain Monte Carlo iterations from the Bayesian implementation of the PTP species delimitation methods (bPTP). Chain does not stay at high likelihood locations but oscillate from high to low locations indicating lack of convergence.

Fig. S2. Relative abundance of the six most abundant bacteria phyla. Each horizontal bar represents a lichen sample and colors reflect different phyla. A. Reindeer lichens samples grouped by molecular species delimitation (multi-rate Poisson tree processes method, mPTP) (host-species dataset). B. *Cladonia stellaris* samples grouped by geography, lichen woodlands northern or southern Quebec (LWs dataset).

Fig. S3. Principal Coordinates analysis (PCoA) of bacterial community composition based on amplicon sequence variants (ASVs) (A) in reindeer lichens, and (B) in *Cladonia stellaris* from lichen woodlands (LWs) northern and southern Quebec. Bacterial communities are not grouped by host species. Two clusters are differentiated (p-value < 0.01) between northern and southern LWs.

Fig. S4. The estimates of richness values (number of observed amplicon sequence variants, ASVs) and alpha diversity indices (Shannon and Simpson) for reindeer lichens bacterial communities. Species delimited by morphology (host-species dataset).

Fig. S5. Relative abundance of the bacteria associated with *Cladonia stellaris*. (A) Bacterial genera with significant (padj = 0.01) log2 fold changes between lichen woodlands (LWs) northern (Kuujjuarapik) and southern (PNGJ) Quebec. “NA” corresponds to amplicon sequence variants (ASVs) whose genera could not be assigned. Each dot represents an ASV. (B) Comparison of bacteria relative abundance between LWs. Groups that significantly differ between Kuujjuarapik and PNGJ are shown, such as genera *Endobacter*, genera *Tundrisphaera*, family Caulobacteraceae and family Emeriobacterota.

Fig. S6. Venn diagram demonstrated the overlaps of the common core bacteriota of reindeer lichens and *Cladonia stellaris*.

Table S1. List of the 192 reindeer lichen specimens collected in Eastern North America and included in the study. The following information is provided for each sample: collection number, species name based on morphological delimitation, species named based on molecular delimitation (multi-rate Poisson tree processes method, mPTP), locality, vegetation zone, bioclimatic domain, altitude, type of genetic data generated and GenBank accession number. Data relative to the twenty-two specimens from GenBank are also provided.

Table S2. Sequencing reads identified among the 189 reindeer lichen specimens before and after quality trimming.

Table S3. Number of amplicon sequence variants (ASVs) for bacterial phyla detected in the two datasets (host-selectivity and LWs). Number of samples and names of reindeer lichen species considered for each dataset are included. Number of ASVs significant different in abundance is also provided.

Table S4. Amplicon sequence variants (ASVs) that significantly differ in relative abundance between (A) reindeer lichen species delimited by the multi-rate Poisson tree processes method (mPTP), (B) reindeer lichen species delimited by morphology, and (C) lichen woodlands (LWs) northern and southern Quebec. P-values and bacteria identity are provided for each ASV. The identity (≥95%) of each ASV according to Blast alignment to 16 rRNA sequences from NCBI database is also included with maximum and total score, query cover and E values.

Table S5. Bacteria amplicon sequence variants (ASVs) with prevalence higher than (A) 0.50 and (B) 0.75 across our reindeer lichen samples (common core bacteriota). For each ASV, we provided phylum, class, order, family, and genus, as well as DNA sequence. The identity (>96%) of each ASV according to Blast alignment to 16 rRNA sequences from NCBI database is also included with maximum and total score, query cover and E values.

Table S6. Total common core abundance in each lichen sample as a sum of abundance of the common core members. A. Data for prevalence of 0.50. B. Data for prevalence of 0.75.

Table S7. Bacterial amplicon sequence variants (ASVs) with prevalence higher than (A) 0.50 and (B) 0.75 across our *Cladonia stellaris* samples (common core bacteriota). For each ASV, we provided Phylum, class, order, family, and genus, as well as DNA sequence. The identity (>96% and >97%) of each ASV according to Blast alignment to 16 rRNA sequences from NCBI database is also included with maximum and total score, query cover and E values.

## Appendix 01

FASTA alignment of 126 ITS sequences belonging to four species of reindeer lichens, such as *Cladonia mitis, C. rangiferina, C. stellaris* and *C. stygia*. Two individuals of *C. wainioi* are included as outgroup.

## Appendix 02

Output file from the multi-rate PTP (mPTP) species delimitation method. Ten species are recognized with strongly supported values. *Cladonia mitis* was divided into four taxa; *C. stellaris* split into three; individuals belonging to *C. rangiferina* and *C. stygia* were merged together but split into two taxa, and two individuals of *C. wainioi* constituted the outgroup.

## Appendix 03

Statistical results of alpha-diversity in reindeer lichens for molecular and morphological-delimitated species (host-selectivity dataset). Values estimated based on number of observed amplicon sequence variants (ASVs), Shannon and Simpson effective indices. Results derived from Shapiro-Wilk test of normality, Kruskal-Wallis non-parametric test and pairwise comparison with U-Mann-Whitney test are shown.

## Appendix 04

Statistical results of beta-diversity in reindeer lichens for molecular and morphological-delimitated species (host-selectivity dataset). Significant differences estimated based on the UniFrac distance matrix. Results derived from PERMANOVA test Adonis and pairwise comparison (pairwise.adonis) are shown.

## Appendix 05

Statistical results of alpha-diversity in *Cladonia stellaris* from lichen woodlands northern and southern Quebec (LWs dataset). Values estimated based on number of observed amplicon sequence variants (ASVs), Shannon and Simpson effective indices. Results derived from Shapiro-Wilk test of normality, Kruskal-Wallis non-parametric test and pairwise comparison with U-Mann-Whitney test are shown.

## Appendix 06

Statistical results of beta-diversity in *Cladonia stellaris* from lichen woodlands (LWs) northern and southern Quebec (LWs dataset). Significant differences estimated based on the UniFrac distance matrix. Results derived from PERMANOVA test Adonis and pairwise comparison (pairwise.adonis) are shown.

## Appendix 07

Results derived from the Blast alignment to sequences in NCBI database 16S ribosomal RNA. Amplicon sequence variants (ASVs) associated to *Cladonia stellaris* significantly different in abundance between northern and southern lichen woodlands were aligned. Sequence identity, maximal and total score, query cover, E values and percentage of identity are displayed.

## Appendix 08

Results derived from the Blast alignment to sequences in NCBI database 16S ribosomal RNA. Amplicon sequence variants (ASVs) included in the common core bacteriota (prevalence 0.50) of reindeer lichens were aligned. Sequence identity, maximal and total score, query cover, E values and percentage of identity are displayed.

## Appendix 09

Results derived from the Blast alignment to sequences in NCBI database 16S ribosomal RNA. Amplicon sequence variants (ASVs) included in the common core bacteriota (prevalence 0.50) of *Cladonia stellaris* were aligned. Sequence identity, maximal and total score, query cover, E values and percentage of identity are displayed.

## Notes

### Competing Interest Statement

The authors have declared no competing interest.

## LITERATURE CITED

ACIA Impacts of a Warming Arctic. 2004. Cambrige, UK: Cambridge University Press.

Agler MT, Ruhe J, Kroll S, et al. 2016. Microbial Hub Taxa Link Host and Abiotic Factors to Plant Microbiome Variation. PLoS Biology 14: e1002352.

Ahti T. 1961. Taxonomic studies on reindeer lichens (Cladonia, subgenus Cladina). Helsinki: Societas Zoologica Botanica Fennica “Vanamo.”

Ainsworth TD, Krause L, Bridge T, et al. 2015. The coral core microbiome identifies rare bacterial taxa as ubiquitous endosymbionts. ISME Journal 9: 2261–2274.

Alonso-García M, Grewe F, Payette S, Villarreal A. JC. 2021. Population genomics of a reindeer lichen species from North-American lichen woodlands. American Journal of Botany.

Aschenbrenner IA, Cardinale M, Berg G, Grube M. 2014. Microbial cargo: Do bacteria on symbiotic propagules reinforce the microbiome of lichens? Environmental Microbiology 16: 3743–3752.

Aschenbrenner IA, Cernava T, Berg G, Grube M. 2016. Understanding Microbial Multi-Species Symbioses. Frontiers in Microbiology 7: 180.

Athukorala SNP, Pino-Bodas R, Stenroos S, Ahti T, Piercey-Normore MD. 2016. Phylogenetic relationships among reindeer lichens of North America. Lichenologist 48: 209–227.

Auclair AND, Rencz AN. 1982. Concentration, mass, and distribution of nutrients in a subarctic Picea mariana - Cladonia alpestris ecosystem (Canada). Canadian Journal of Forest Research 12: 947–968.

Bates ST, Cropsey GWG, Caporaso JG, Knight R, Fierer N. 2011. Bacterial communities associated with the lichen symbiosis. Applied and Environmental Microbiology 77: 1309–1314.

Belova SE, Kulichevskaya IS, Bodelier PLE, Dedysh SN. 2013. Methylocystis bryophila sp. nov., a facultatively methanotrophic bacterium from acidic Sphagnum peat, and emended description of the genus Methylocystis (ex Whittenbury et al. 1970) Bowman et al. 1993. International Journal of Systematic and Evolutionary Microbiology 63: 1096–1104.

Berestovskaya JJ, Kotsyurbenko OR, Tourova TP, et al. 2012. Methylorosula polaris gen. nov., sp. nov., an aerobic, facultatively methylotrophic psychrotolerant bacterium from tundra wetland soil. International Journal of Systematic and Evolutionary Microbiology 62: 638–646.

Berg G, Rybakova D, Fischer D, et al. 2020. Microbiome definition re-visited: old concepts and new challenges. Microbiome 8: 103.

Blair C, Bryson RW. 2017. Cryptic diversity and discordance in single-locus species delimitation methods within horned lizards (Phrynosomatidae: Phrynosoma). Molecular Ecology Resources 17: 1168–1182.

Bouchard R, Peñaloza-Bojacá G, Toupin S, et al. 2020. Contrasting bacteriome of the hornwort Leiosporoceros dussii in two nearby sites with emphasis on the hornwort-cyanobacterial symbiosis. Symbiosis 81: 39–52.

Bragina A, Berg C, Berg G. 2015. The core microbiome bonds the Alpine bog vegetation to a transkingdom metacommunity. Molecular Ecology 24: 4795–4807.

Brodo IM, Sharnoff SD, Sharnoff S. 2001. Lichens of North America. New Haven: Yale University Press.

Bubrick P, Frensdorff A, Galun M. 1985. Selectivity in the Lichen Symbiosis In: Lichen Physiology and Cell Biology. Springer US, 319–334.

Callahan BJ, McMurdie PJ, Holmes SP. 2017. Exact sequence variants should replace operational taxonomic units in marker-gene data analysis. ISME Journal 11: 2639–2643.

Callahan BJ, Sankaran K, Fukuyama JA, McMurdie PJ, Holmes SP. 2016. Bioconductor workflow for microbiome data analysis: From raw reads to community analyses [version 1; referees: 3 approved]. F1000Research 5: 1–49.

Cardinale M, Grube M, Vieirade Castro Jr J, Müller H, Berg G. 2012. Bacterial taxa associated with the lung lichen Lobaria pulmonaria are differentially shaped by geography and habitat. FEMS Microbiology Letters: 111–115.

Cardinale M, Steinová J, Rabensteiner J, Berg G, Grube M. 2012. Age, sun and substrate: Triggers of bacterial communities in lichens. Environmental Microbiology Reports 4: 23–28.

Cardinale M, Vieira de Castro J, Muller H, Berg G, Grube M. 2008. In situ analysis of the bacterial community associated with the reindeer lichen Cladonia arbuscula reveals predominance of Alphaproteobacteria. FEMS Microbiology Ecology 66: 63–71.

Cernava T, Erlacher A, Aschenbrenner IA, et al. 2017. Deciphering functional diversification within the lichen microbiota by meta-omics. Microbiome 5: 82.

Chappell TM, Rausher MD. 2016. Evolution of host range in Coleosporium ipomoeae, a plant pathogen with multiple hosts. Proceedings of the National Academy of Sciences of the United States of America 113: 5346–5351.

Coyte KZ, Rakoff-Nahoum S. 2019. Understanding Competition and Cooperation within the Mammalian Gut Microbiome. Current Biology 29: R538–R544.

Cullen CM, Aneja KK, Beyhan S, et al. 2020. Emerging Priorities for Microbiome Research. Frontiers in Microbiology 11: 136.

David LA, Maurice CF, Carmody RN, et al. 2014. Diet rapidly and reproducibly alters the human gut microbiome. Nature 505: 559–563.

Delgado-Baquerizo M, Oliverio AM, Brewer TE, et al. 2018. A global atlas of the dominant bacteria found in soil. Science 359: 320–325.

Dellicour S, Flot JF. 2018. The hitchhiker’s guide to single-locus species delimitation. Molecular Ecology Resources 18: 1234–1246.

Denison ER, Rhodes RG, McLellan WA, Pabst DA, Erwin PM. 2020. Host phylogeny and life history stage shape the gut microbiome in dwarf (Kogia sima) and pygmy (Kogia breviceps) sperm whales. Scientific Reports 10: 1–13.

Douglas AE. 2018. The Drosophila model for microbiome research. Lab Animal 47: 157–164.

Erlacher A, Cernava T, Cardinale M, et al. 2015. Rhizobiales as functional and endosymbiontic members in the lichen symbiosis of Lobaria pulmonaria L. Frontiers in Microbiology 6: 53.

Faure D, Simon JC, Heulin T. 2018. Holobiont: a conceptual framework to explore the eco-evolutionary and functional implications of host–microbiota interactions in all ecosystems. New Phytologist 218: 1321–1324.

Federici S, Nobs SP, Elinav E. 2020. Phages and their potential to modulate the microbiome and immunity. Cellular and Molecular Immunology: 1–16.

Fernández-Brime S, Muggia L, Maier S, Grube M, Wedin M. 2019. Bacterial communities in an optional lichen symbiosis are determined by substrate, not algal photobionts. FEMS Microbiology Ecology 95: 1–11.

Fuentes-Ramírez LE, Bustillos-Cristales R, Tapia-Hernández A, et al. 2001. Novel nitrogen-fixing acetic acid bacteria, Gluconacetobacter johannae sp. nov. and Gluconacetobacter azotocaptans sp. nov., associated with coffee plants. International Journal of Systematic and Evolutionary Microbiology 51: 1305–1314.

Grasby SE, Richards BC, Sharp CE, Brady AL, Jones GM, Dunfield PF. 2013. The paint pots, Kootenay National Park, Canada - a natural acid spring analogue for mars. Canadian Journal of Earth Sciences 50: 94–108.

Greenberg DE, Porcella SF, Stock F, et al. 2006. Granulibacter bethesdensis gen. nov., sp. nov., a distinctive pathogenic acetic acid bacterium in the family Acetobacteraceae. International Journal of Systematic and Evolutionary Microbiology 56: 2609–2616.

Grube M, Berg G. 2009. Microbial consortia of bacteria and fungi with focus on the lichen symbiosis. Fungal Biology Reviews 23: 72–85.

Grube M, Cardinale M, De Castro JV, Müller H, Berg G. 2009. Species-specific structural and functional diversity of bacterial communities in lichen symbioses. ISME Journal 3: 1105–1115.

Grube M, Cernava T, Soh J, et al. 2015. Exploring functional contexts of symbiotic sustain within lichen-associated bacteria by comparative omics. ISME Journal 9: 412–424.

Hasegawa M, Kishino H, Yano T aki. 1985. Dating of the human-ape splitting by a molecular clock of mitochondrial DNA. Journal of Molecular Evolution 22: 160–174.

Hassani MA, Durán P, Hacquard S. 2018. Microbial interactions within the plant holobiont. Microbiome 6: 58.

Hodkinson BP, Gottel NR, Schadt CW, Lutzoni F. 2012. Photoautotrophic symbiont and geography are major factors affecting highly structured and diverse bacterial communities in the lichen microbiome. Environmental Microbiology 14: 147–161.

Hodkinson BP, Lutzoni F. 2009. A microbiotic survey of lichen-associated bacteria reveals a new lineage from the Rhizobiales. Symbiosis 49: 163–180.

Holland-Moritz H, Stuart J, Lewis LR, et al. 2018. Novel bacterial lineages associated with boreal moss species. Environmental Microbiology 20: 2625–2638.

Huse SM, Ye Y, Zhou Y, Fodor AA. 2012. A Core Human Microbiome as Viewed through 16S rRNA Sequence Clusters (N Ahmed, Ed.). PLoS ONE 7: e34242.

Ivanova AA, Kulichevskaya IS, Merkel AY, Toshchakov S V., Dedysh SN. 2016. High Diversity of Planctomycetes in Soils of Two Lichen-Dominated Sub-Arctic Ecosystems of Northwestern Siberia. Frontiers in Microbiology 7: 2065.

Jasinski JPP, Payette S. 2005. The creation of alternative stable states in the southern boreal forest, Québec, Canada. Ecological Monographs 75: 561–583.

Jobbágy EG, Jackson RB. 2000. The vertical distribution of soil organic carbon and its relation to climate and vegetation. Ecological Applications 10: 423–436.

Johnson EA, Miyanishi K. 1999. Subarctic lichen woodlands In: Anderson R, Fralish J, Baskin J, eds. Savanna, barren and rock outcrop plant communities of North America. Cambrige, UK: Cambridge University Press, 421–436.

Jorge F, Dheilly NM, Poulin R. 2020. Persistence of a Core Microbiome Through the Ontogeny of a Multi-Host Parasite. Frontiers in Microbiology 11.

Kapli P, Lutteropp S, Zhang J, et al. 2017. Multi-rate Poisson tree processes for single-locus species delimitation under maximum likelihood and Markov chain Monte Carlo. Bioinformatics 33: 1630–1638.

Kistenich S, Bendiksby M, Ekman S, Cáceres MES, Hernández JEM, Timdal E. 2019. Towards an integrative taxonomy of Phyllopsora (Ramalinaceae). Lichenologist 51: 323–392.

Kulichevskaya IS, Ivanova AA, Detkova EN, et al. 2017. Tundrisphaera lichenicola gen. nov., sp. nov., a psychrotolerant representative of the family Isosphaeraceae from lichen-dominated tundra soils. International Journal of Systematic and Evolutionary Microbiology 67: 3583–3589.

Lahti L. Tools for microbiome analysis in R. Microbiome package version 4.

Larsen. 2014. Impacts, Adaptation, and Vulnerability In: Field CB, ed. Climatic Change. Cambrige, NY: Cambridge University Press, 1567–1612

Lavoie C, Renaudin M, McMullin RT, et al. 2020. Extremely low genetic diversity of Stigonema associated with Stereocaulon in eastern Canada. The Bryologist 123: 188.

Lee MD, Kling JD, Araya R, Ceh J. 2018. Jellyfish Life Stages Shape Associated Microbial Communities, While a Core Microbiome Is Maintained Across All. Frontiers in Microbiology 9: 1534.

Ley RE, Hamady M, Lozupone C, et al. 2008. Evolution of mammals and their gut microbes. Science 320: 1647–1651.

Lloyd-Price J, Mahurkar A, Rahnavard G, et al. 2017. Strains, functions and dynamics in the expanded Human Microbiome Project. Nature 550: 61–66.

Love MI, Huber W, Anders S. 2014. Moderated estimation of fold change and dispersion for RNA-seq data with DESeq2. Genome Biology 15: 550.

Margulis L. 1991. Symbiosis as a source of evolutionary innovation: speciation and morphogenesis In: MA, MLFR, eds. Symbiogenesis and Symbionticism. Cambrige, NY: MIT Press, 1–14

Marki PZ, Fjeldså J, Irestedt M, Jønsson KA. 2018. Molecular phylogenetics and species limits in a cryptically coloured radiation of Australo-Papuan passerine birds (Pachycephalidae: Colluricincla). Molecular Phylogenetics and Evolution 124:100–105

Marks RA, Smith JJ, Cronk Q, McLetchie DN. 2017. Variation in the bacteriome of the tropical liverwort, Marchantia inflexa, between the sexes and across habitats. Symbiosis 75: 93–101.

Martinez Arbizu P. 2020. pairwiseAdonis: Pairwise multilevel comparison using adonis. R package version 0.4.

McFall-Ngai M. 2008. Are biologists in “future shock”? Symbiosis integrates biology across domains. Nature Reviews Microbiology 6: 789–792.

McLaren MR. 2020. Silva SSU taxonomic training data formatted for DADA2 (Silva version 138).

McMurdie PJ, Holmes S. 2013. phyloseq: An R Package for Reproducible Interactive Analysis and Graphics of Microbiome Census Data (M Watson, Ed.). PLoS ONE 8: e61217.

Morneau C, Payette S. 1989. Postfire lichen-spruce woodland recovery at the lmit of the boreal forest in northern Quebec. Canadian Journal of Botany 67: 2770–2782.

Morrissey KL, Çavaş L, Willems A, De Clerck O. 2019. Disentangling the Influence of Environment, Host Specificity and Thallus Differentiation on Bacterial Communities in Siphonous Green Seaweeds. Frontiers in Microbiology 10: 717.

Mushegian AA, Peterson CN, Baker CCM, Pringle A. 2011. Bacterial diversity across individual lichens. Applied and Environmental Microbiology 77: 4249–4252.

Nishijima M, Tazato N, Handa Y, et al. 2013. Gluconacetobacter tumulisoli sp. nov., Gluconacetobacter Takamatsuzukensis sp. nov. and Gluconacetobacter aggeris sp. nov., isolated from Takamatsuzuka tumulus samples before and during the dismantling work in 2007. International Journal of Systematic and Evolutionary Microbiology 63: 3981–3988.

Noh H-J, Lee YM, Park CH, Lee HK, Cho J-C, Hong SG. 2020. Microbiome in Cladonia squamosa Is Vertically Stratified According to Microclimatic Conditions. Frontiers in Microbiology 11.

Nylander JAA. 2004. MrModeltest Version 2. Program Distributed by the Author.

Oksanen J, Blanchet FG, Friendly M, et al. 2020. Vegan: community ecology package. http://cran.r-project.org/package=vegan. 30 Nov. 2020.

Pan X, Yang Y, Zhang J-R. 2014. Molecular basis of host specificity in human pathogenic bacteria. Emerging Microbes & Infections 3: 1–10.

Park S, Jang S, Oh S, Kim JA, Hur J. 2014. An Easy, Rapid, and Cost-Effective Method for DNA Extraction from Various Lichen Taxa and Specimens Suitable for Analysis of Fungal and Algal Strains. Mycobiology 42: 311–320.

Park CH, Kim KM, Kim OS, Jeong G, Hong SG. 2016. Bacterial communities in Antarctic lichens. Antarctic Science 28: 455–461.

Payette S. 1992. Fire as a controlling process in the North American boreal forest In: Herman H. S, Rik L, Gordon B. B, eds. A systems analysis of the global boreal forest. Cambrige, NY: Cambridge University Press, 144–169

Pearce DS, Hoover BA, Jennings S, Nevitt GA, Docherty KM. 2017. Morphological and genetic factors shape the microbiome of a seabird species (Oceanodroma leucorhoa) more than environmental and social factors. Microbiome 5: 146.

Pr Herlemann D, Labrenz M, Jü Rgens K, Bertilsson S, Waniek JJ, Andersson AF. 2011. Transitions in bacterial communities along the 2000 km salinity gradient of the Baltic Sea. The ISME Journal 5: 1571–1579.

Printzen C, Fernández-Mendoza F, Muggia L, Berg G, Grube M. 2012. Alphaproteobacterial communities in geographically distant populations of the lichen *Cetraria aculeata*. FEMS Microbiology Ecology 82: 316–325.

Rahme LG, Ausubel FM, Cao H, et al. 2000. Plants and animals share functionally common bacterial virulence factors. Proceedings of the National Academy of Sciences of the United States of America 97: 8815–8821.

Relman DA. 2008. “Til death do us part”: Coming to terms with symbiotic relationships. Nature Reviews Microbiology 6: 721–724.

Reveillaud J, Maignien L, Eren MA, et al. 2014. Host-specificity among abundant and rare taxa in the sponge microbiome. ISME Journal 8: 1198–1209.

Risely A. 2020. Applying the core microbiome to understand host–microbe systems (A Tate, Ed.). Journal of Animal Ecology 89: 1549–1558.

Roi GHL. 2018. Boreal Zone. The Canadian Encyclopedia.

Ronquist F, Teslenko M, Van Der Mark P, et al. 2012. MrBayes 3.2: Efficient Bayesian Phylogenetic Inference and Model Choice Across a Large Model Space. Systematic Biology 61: 539–542.

Rothschild D, Weissbrod O, Barkan E, et al. 2018. Environment dominates over host genetics in shaping human gut microbiota. Nature 555: 210–215.

Rowe JS. 1972. Forest Regions of Canada. Ottawa: Canadian Forest Service Publications 1300.

Saravanan VS, Madhaiyan M, Osborne J, Thangaraju M, Sa TM. 2008. Ecological occurrence of Gluconacetobacter diazotrophicus and nitrogen-fixing Acetobacteraceae members: Their possible role in plant growth promotion. Microbial Ecology 55: 130–140.

Schlechter RO, Miebach M, Remus-Emsermann MNP. 2019. Driving factors of epiphytic bacterial communities: A review. Journal of Advanced Research 19: 57–65.

Schliep KP. 2011. phangorn: Phylogenetic analysis in R. Bioinformatics 27: 592–593.

Schliep K, Potts AJ, Morrison DA, Grimm GW. 2017. Intertwining phylogenetic trees and networks (R Fitzjohn, Ed.). Methods in Ecology and Evolution 8: 1212–1220.

Sepulveda J, Moeller AH. 2020. The Effects of Temperature on Animal Gut Microbiomes. Frontiers in Microbiology 11: 384.

Shaver GR, Chapin FS. 1991. Production: Biomass relationships and element cycling in contrasting arctic vegetation types. Ecological Monographs 61: 1–23.

Sierra MA, Danko DC, Sandoval TA, et al. 2020. The Microbiomes of Seven Lichen Genera Reveal Host Specificity, a Reduced Core Community and Potential as Source of Antimicrobials. Frontiers in Microbiology 11: 398.

Sigurbjörnsdóttir MA, Andrésson S. Ó, Vilhelmsson O. 2015. Analysis of the Peltigera membranacea metagenome indicates that lichen-associated bacteria are involved in phosphate solubilization. Microbiology 161: 989–996.

Simon JC, Marchesi JR, Mougel C, Selosse MA. 2019. Host-microbiota interactions: From holobiont theory to analysis. Microbiome 7: 5.

Skogland T. 1984. Wild reindeer foraging-niche organization. Ecography 7: 345–379.

Stenroos S, Pino-Bodas R, Hyvönen J, Lumbsch HT, Ahti T. 2018. Phylogeny of the family Cladoniaceae (Lecanoromycetes, Ascomycota) based on sequences of multiple loci. Cladistics 35: 351–384.

Svihus B, Holand Ø. 2000. Lichen polysaccharides and their relation to reindeer/caribou `nutrition`.

Tarnocai C, Canadell JG, Schuur EAG, Kuhry P, Mazhitova G, Zimov S. 2009. Soil organic carbon pools in the northern circumpolar permafrost region. Global Biogeochemical Cycles 23: 1–11.

Thompson ID, Wiebe PA, Mallon E, et al. 2015. Factors influencing the seasonal diet selection by woodland caribou (rangifer tarandus tarandus) in boreal forests in ontario. Canadian Journal of Zoology 93: 87–98.

Trexler R, Solomon C, Brislawn CJ, et al. 2014. Assessing impacts of unconventional natural gas extraction on microbial communities in headwater stream ecosystems in Northwestern Pennsylvania. Frontiers in Microbiology 5: 522.

Trivedi P, Leach JE, Tringe SG, Sa T, Singh BK. 2020. Plant-microbiome interactions: from community assembly to plant health Nature reviews | Microbiology. Nature Reviews Microbiology.

Turnbaugh PJ, Ley RE, Hamady M, Fraser-Liggett CM, Knight R, Gordon JI. 2007. The Human Microbiome Project. Nature 449: 804–810.

Vandenkoornhuyse P, Quaiser A, Duhamel M, Le Van A, Dufresne A. 2015. The importance of the microbiome of the plant holobiont. New Phytologist 206: 1196–1206.

Vincent AT, Derome N, Boyle B, Culley AI, Charette SJ. 2017. Next-generation sequencing (NGS) in the microbiological world: How to make the most of your money. Journal of Microbiological Methods 138: 60–71.

Wagner MR, Lundberg DS, Del Rio TG, Tringe SG, Dangl JL, Mitchell-Olds T. 2016. Host genotype and age shape the leaf and root microbiomes of a wild perennial plant. Nature Communications 7: 12151.

Youngblut ND, Reischer GH, Walters W, et al. 2019. Host diet and evolutionary history explain different aspects of gut microbiome diversity among vertebrate clades. Nature Communications 10: 3500.

Zhang J, Kapli P, Pavlidis P, Stamatakis A. 2013. A general species delimitation method with applications to phylogenetic placements. Bioinformatics 29: 2869–2876.

Zheng Y, Gong X. 2019. Niche differentiation rather than biogeography shapes the diversity and composition of microbiome of Cycas panzhihuaensis. Microbiome 7: 152.

